# Gene Expression Meta-Analysis Reveals Interferon-induced Genes

**DOI:** 10.1101/2020.11.14.382697

**Authors:** Amber Park, Laura K. Harris

## Abstract

**Background:** Severe Acute Respiratory Syndrome (SARS) corona virus (CoV) infections are a serious public health threat because of their pandemic-causing potential. This work analyzes mRNA expression data from SARS infections through meta-analysis of gene signatures, possibly identifying therapeutic targets associated with major SARS infections.

**Methods:** This work defines 37 gene signatures representing SARS-CoV, Middle East Respiratory Syndrome (MERS)-CoV, and SARS-CoV2 infections in human lung cultures and/or mouse lung cultures or samples and compares them through Gene Set Enrichment Analysis (GSEA). To do this, positive and negative infectious clone SARS (icSARS) gene panels are defined from GSEA-identified leading-edge genes between two icSARS-CoV derived signatures, both from human cultures. GSEA then is used to assess enrichment and identify leading-edge icSARS panel genes between icSARS gene panels and 27 other SARS-CoV gene signatures. The meta-analysis is expanded to include five MERS-CoV and three SARS-CoV2 gene signatures. Genes associated with SARS infection are predicted by examining the intersecting membership of GSEA-identified leading-edges across gene signatures.

**Results:** Significant enrichment (GSEA p<0.001) is observed between two icSARS-CoV derived signatures, and those leading-edge genes defined the positive (233 genes) and negative (114 genes) icSARS panels. Non-random significant enrichment (null distribution p<0.001) is observed between icSARS panels and all verification icSARSvsmock signatures derived from human cultures, from which 51 over- and 22 under-expressed genes are shared across leading-edges with 10 over-expressed genes already associated with icSARS infection. For the icSARSvsmock mouse signature, significant, non-random significant enrichment held for only the positive icSARS panel, from which nine genes are shared with icSARS infection in human cultures. Considering other SARS strains, significant, non-random enrichment (p<0.05) is observed across signatures derived from other SARS strains for the positive icSARS panel. Five positive icSARS panel genes, CXCL10, OAS3, OASL, IFIT3, and XAF1, are found across mice and human signatures regardless of SARS strains.

**Conclusion:** The GSEA-based meta-analysis approach used here identifies genes with and without reported associations with SARS-CoV infections, highlighting this approach’s predictability and usefulness in identifying genes that have potential as therapeutic targets to preclude or overcome SARS infections.

## 1 Background

Human β-coronaviruses (CoV) are enveloped, positive-sense RNA viruses that infect humans and a variety of animal species [1]. Human CoV infections typically cause mild upper respiratory distress, referred to as the common cold, and generally were non-lethal [1, 2]. However, in 2002 a novel CoV was found to cause the potentially life-threatening disease, severe acute respiratory syndrome (SARS). The initial SARS-CoV outbreak infected an estimated 8,400 people with over a 9% mortality rate [2–5]. In 2012, another highly lethal CoV causing Middle East Respiratory Syndrome (MERS) emerged with an over 30% mortality rate [6, 7]. Fortunately, both the SARS-CoV and MERS-CoV outbreaks were quickly contained through aggressive infection control measures. Efforts are still ongoing to control the pandemic of SARS-CoV2, a closely related CoV strain with almost 80% genome similarity to SARS-CoV and 50% similarity to MERS-CoV [8, 9]. SARS-CoV2 is the causative agent of coronavirus disease 2019 (COVID-19), which is already responsible for over a 3.7 million deaths worldwide as of June 4 2021 [10–12]. Several new variants of SARS-CoV2 are circulating the globe already, with some variants having increased infectiousness such as B.1.1.7 which originated in the United Kingdom in early 2021, prompting scientists to consider outbreaks of a future SARS-CoV3 strain [10, 13–15].

Options to successfully treat SARS-infected patients are critical to improving patient outcomes. Limited therapeutic options for SARS-CoV infections were available at the time. For example, the RNA-dependent RNA polymerase inhibitor, ribavirin, and corticosteroids were cornerstones of SARS-CoV treatment. Unfortunately, several reports exist showing a lack of efficacy for ribavirin and debating the effectiveness of corticosteroid therapy in SARS-CoV treatment [5, 16]. Remdesivir is a nucleotide analogue that inhibits viral RNA synthesis in SARS via an unknown mechanism. Remdesivir achieved mixed results when treating SARS patients [17–19]. MERS-CoV treatment options were limited also, with most therapies being existing drugs for other disease indications shown to target MERS-CoV replication *in vitro* [20]. Clinical therapies previously used to treat SARS-CoV, like ribavirin and remdesivir, were common treatments for MERS-CoV infections and there was little investment in developing new therapeutic options for SARS infections [21]. That initiative changed dramatically over the past year with many resources devoted to improving treatment options for SARS-CoV2 due to its pandemic status. For example, over 284 clinical trials at different phases are underway to examine efficacy and safety for repurposed drug and new therapeutic molecule development as potential treatment options against COVID-19 [21]. Therapies previously used to treat SARS-CoV and MERS-CoV, like ribavirin and remdesivir, are being re-evaluated as treatments for COVID-19 [18, 19, 21]. Newer therapies targeting immune response, particularly the cytokine storm described frequently in COVID-19 patients, such as interferon (IFN) and interleukin (IL)-6 receptor antagonist therapies, are emerging prominently as clinical options [21–27]. Inhibitors of membrane fusion, viral replicases, and human and viral proteases also are under extensive examination as potential therapeutics against COVID-19 [21]. While these various treatment options show promise, no specific therapy is available currently to treat SARS-CoV2 infections as death tolls from the virus continue to increase [24]. Fortunately, several SARS-CoV2 vaccines are available now as a preventative measure [28, 29]. New therapeutic strategies are needed to successfully treat current and future SARS infections.

A complete understanding of the molecular changes driving SARS infections can assist in the development of new therapies to fight COVID-19 and other future SARS outbreaks. Many studies have been done to elucidate molecular changes associated with a SARS infection by examining gene expression changes and some changes have been confirmed via experimental techniques such as quantitative real-time polymerase chain reaction (qRT-PCR). These studies conducted differential expression analysis of individual genes using statistical methods such as fold change and/or T-test p-value to identify genes of interest [2, 30–34]. This approach usually generates an exhaustive list of several hundred genes that meet pre-established statistical criteria (*e.g.*, fold change <2 and/or T-test p-value<0.05). Further analysis is needed to refine lists of identified genes to identify the most promising gene candidates to target therapeutically. Since experimental work is time consuming and expensive, computational approaches to interpret and prioritize gene lists are used widely.

Several different computational approaches have been used to interpret experimentally generated differentially expressed gene lists. For example, functional enrichment was performed to interpret generated gene lists using a variety of bioinformatic approaches including network analysis [30, 31, 34, 35] and/or hypergeometric test-based approaches (*e.g.*, Fisher’s Exact Test) that utilize established gene sets from public knowledgebases like Gene Ontology (GO) [2, 31–33]. These studies identified several gene expression changes associated with SARS-CoV infection including increased expression of inflammatory mediators IL-6, IL-8, CXCL10 (*i.e.*, INF γ-induced protein 10), IFN-λ, IFIT1, OASL, and OAS3 in SARS-CoV infected human lung cultures and mouse lung samples [30, 32, 34, 35]. Comparative analysis of differential gene expression across SARS strains found significant increases in gene expression of some genes, like CXCL10, IFIT1, and OAS3, or their gene relations, such as CXCL1, CXCL2, and IFIT3, in MERS-CoV and/or SARS-CoV2 infections in human epithelial lung cells and lung samples from cynomolgus monkey and mice [36–39]. Other genes were unique to a SARS strain, such as XAF1 which has been reported only in SARS-CoV2 infections [36–39]. Hypergeometric test-based approaches are known to be limited because only genes that meet an established cut-off (*e.g.*, T-test p-value<0.05) are considered [40]. To overcome this limitation by considering all genes in an expression dataset, other studies utilized Gene Set Enrichment Analysis (GSEA) to calculate the enrichment of an established gene set from a public knowledgebase (*e.g.*, MSigDB, Blood Transcriptional Modules, and/or Kyoto Encyclopedia of Genes and Genomes (KEGG) pathways) in a gene signature. This was done by defining a gene list ranked by differential expression between mock and SARS infected samples by an appropriate statistical method [30, 41]. Findings from these studies included positively enriched modules associated with antiviral IFN, cell cycle and proliferation, and monocytes and dendritic cells in peripheral blood mononuclear cells from SARS-CoV2 patients [41]. Further, GSEA has been used to confirm enrichment for a query gene set defined from a network analysis approach, and identified regulatory genes associated with the pathogenicity of SARS-CoV have been identified [30]. This demonstrated that GSEA is a useful computational tool to validate gene candidates, though all reported studies had used GSEA on established query genes sets rather than using GSEA to establish new sets. Therefore, we sought to use a GSEA-based approach to identify gene candidates from SARS mRNA expression datasets here.

In a prior study to identify genes associated with macrolide resistance in *Streptococcus pneumoniae*, we demonstrated a GSEA-approach for gene identification that compared differential gene expression between mRNA expression datasets [42]. This study successfully identified known and novel genes though it was limited due to incomplete genome annotation, a common problem for many microbial genomes [43–50]. Applying our GSEA-approach to mRNA expression data derived from a species with a more completely annotated genome, such as human or mouse, our achievable results would improve. Therefore, in this paper we use the GSEA-approach previously used on *Streptococcus pneumoniae* to identify gene expression changes associated with SARS infection in human lung cell cultures and mouse lung samples. Using GSEA in this way generates a list of gene candidates associated with a SARS infection which is similar in nature to gene candidate lists generated previously by using single-gene analysis. To refine and prioritize the exhaustive gene lists generated from such an analysis, we performed a GSEA-based meta-analysis that incorporates over 35 different gene signatures. We hypothesized that therapeutically targeting gene candidates identified from our nested GSEA meta-analysis has the potential to improve treatment options for SARS infections. However, we acknowledge that this work is entirely computational and further experimental and clinical work will be needed to properly validate our findings and implement them clinically.

## 2 Methods

### 2.1 mRNA Expression Resources

#### 2.1.1 SARS-CoV Datasets

To identify molecular changes associated with SARS infection in human lung cell cultures, we searched the Gene Expression Omnibus (GEO) repository [51–53] to find seven datasets for use in our study (Table 1). Since most datasets contained samples infected with an infectious clone of SARS-CoV Urbani (icSARS), we selected the published SuperSeries GSE47963, which contained three independent datasets (GSE47960, GSE47961, and GSE47962) that examined gene expression in icSARS and mock infected human primary tracheobronchial epithelium cells [30]. We used this SuperSeries to identify and verify gene expression changes associated with icSARS infection mechanisms. These datasets also contained gene expression data for cells infected with two other strains: 1) icSARS-dORF6, the icSARS strain with a genetic deletion causing a lack of expression of ORF6 which encodes an IFN antagonist protein that increases virulence by blocking human nuclear translocation [2, 3, 30], or 2) SARS-BatSRBD, a SARS-CoV like virus isolated from bats that was synthetically modified to contain the spike receptor binding domain (SRBD) from the wild type Urbani strain to allow for infection of human and non-human primate cells [30, 54]. To extend our verification of identified gene expression changes associated with icSARS infection into another lung cell type, we selected unpublished datasets GSE37827 and GSE48142 that examined gene expression in icSARS and mock infected human lung adenocarcinoma 2B-4 cells. GSE37827 also contained SARS-BatSRBD infected samples, and GSE48142 had samples infected with mutant strains containing either 1) a genetic modification of NSP16 (deltaNSP16) that encodes a non-structural 2’O methyltransferase whose disruption increases sensitivity to type 1 and 3 IFN responses [33], or 2) ExoNI, which has no formal description in public knowledgebases. All gene expression data for these five datasets were profiled on the commercial probe name version of the Agilent-014850 Whole Human Genome Microarray 4×44K G4112F platform (GEO GPL6480). Further, we selected GSE33267 because it had mock, icSARS, and dORF6 infected samples of Calu-3 cells, from which 2B-4 was a clonal derivative. GSE33267 was profiled on the feature number version of the Agilent-014850 Whole Human Genome Microarray 4×44K G4112F (GEO GPL4133). To compare identified changes associated with icSARS infection to changes associated with the icSARS’s parent Urbani SARS strain, we selected GSE17400 since it had samples of 2B-4 with mock or Urbani infection [32]. GSE17400 was profiled on the Affymetrix Human Genome U133 Plus 2.0 Array (GEO GPL570).

**Table 1.** SARS-CoV Infection Datasets Utilized for this Study

| Dataset | Description<sup>1</sup> | Platform | Probes | IDs | Genes |
| --- | --- | --- | --- | --- | --- |
| GSE47960 | Unnormalized cultures of human primary tracheobronchial epithelium cells with mock, infectious clone SARS-CoV (icSARS) derived from Urbani strain, SARS-dORF6, or SARS-BatSRBD infection | GPL6480 | 32067 | 25666 | 16479 |
| GSE47961 | Unnormalized cultures of human primary tracheobronchial epithelium cells with mock, icSARS, SARS-dORF6, or SARS-BatSRBD infection | GPL6480 | 33421 | 26500 | 16978 |
| GSE47962 | Unnormalized cultures of human primary tracheobronchial epithelium cells with mock, icSARS, SARS-dORF6, or SARS-BatSRBD infection | GPL6480 | 32388 | 25897 | 16603 |
| GSE37827 | Unnormalized cultures of 2B4 cells (a clonal derivative of human lung adenocarcinoma Calu-3 cells) with mock, icSARS, or SARS-BatSRBD infection | GPL6480 | 41000 | 30936 | 19596 |
| GSE48142 | Unnormalized cultures of human lung adenocarcinoma 2B4 cells with mock, icSARS, icSARS-ExoNI or icSARS-dNSP16 infection | GPL6480 | 33321 | 26212 | 16794 |
| GSE33267 | Unnormalized cultures of human lung adenocarcinoma Calu3cells with mock, icSARS, or SARS-dORF6 infection | GPL4133 | 45015 | 32696 | 19751 |
| GSE17400 | Unnormalized cultures of human lung adenocarcinoma 2B4 cells with mock or Urbani strain of SARS-CoV (SARS) infection | GPL570 | 54675 | 44134 | 21182 |
| GSE50000 | Unnormalized lung samples from 20-week-old C57BL6 mice infected with mock, icSARS, either 10<sup>4</sup> or 10<sup>5</sup> plaque-forming unit wild-type mouse-adapted SARS MA15 or SARS-BatSRBD | GPL7202 | 41174 | 34907 | 21200 |
| GSE33266 | Unnormalized lung samples from 20-week-old C57BL6 mice infected with mock or wild-type mouse-adapted SARS MA15 at 10<sup>2</sup>, 10<sup>3</sup>, 10<sup>4</sup>, or 10<sup>5</sup> plaque-forming units over a time course | GPL4134 | 45057 | 36933 | 21608 |
| GSE49262 | Log2 normalized lung samples from 20-week-old C57BL6 mice infected with mock, 10<sup>5</sup> plaque-forming unit wild-type SARS MA15, or SARS-dORF6 | GPL7202 | 41174 | 34907 | 21200 |
| GSE49263 | Log2 normalized lung samples from 20-week-old C57BL6 mice infected with mock, 10<sup>5</sup> plaque-forming unit wild-type SARS MA15, or SARS-dNSP16 | GPL7202 | 41174 | 34907 | 21200 |

<sup>1</sup>All datasets collected samples over a time course. 48hr samples were used for this study unless otherwise stated. Unnormalized samples were z-scored across all dataset samples regardless of time or infection for normalization before use. Platform, Gene Expression Omnibus platform identifiers. Probes, number of platform probes with expression data in dataset. IDs, number of probes with Gene Expression Omnibus provided Entrez identifiers. Genes, number of unique Entrez identifiers used for gene signature definition with representatives for duplicate Entrez identifiers being selected by highest coefficient of variation. Values provided in the Genes column reflect the number of genes in gene signatures defined for this study.

To determine if the molecular changes associated with SARS infection observed in human lung cell cultures were reproducible in an *in vivo* mouse model, we selected four datasets that examined lung samples from mock or SARS-CoV infected 20-week-old C57BL6 mice. GSE50000 contained samples from mice infected with icSARS, SARS-BatSRBD, or MA15, a mouse adapted SARS-CoV strain, at two different inoculation doses (10^4^ and 10^5^). GSE33266 had samples over an inoculation dose range (10^2^, 10^3^, 10^4^, and 10^5^). GSE49262 had samples from mice infected with MA15 (10^5^ inoculation dose) or dORF6 mutant strains, and GSE49263 had samples infected with MA15 (10^5^ inoculation dose) or dNSP16 mutant strains. All gene expression data for these datasets were profiled on the Probe Name version of the commercial Agilent-014868 Whole Mouse Genome Microarray 4×44K G4122F platform (GEO GPL7202), except for GSE33266 which was profiled on the Feature Number version of the same platform (GEO GPL4134).

All SARS-CoV datasets measured gene expression over a time course, so we used time-matched samples collected at 48hrs post-infection unless otherwise stated. Expression data provided by GEO for all datasets except GSE49262 and GSE49263 were unnormalized intensities, so we z-scored across all samples within the dataset for normalization prior to use when appropriate. For data cleaning, we converted probes to Entrez ID for each gene using the GEO provided platform data table when necessary. Probes with no provided Entrez ID were removed from analysis. If multiple probes with the same Entrez ID existed, the probe with the highest coefficient of variance across duplicate probes was selected.

#### 2.1.2 MERS-CoV Datasets

To compare molecular changes identified in SARS-CoV to changes associated with MERS-CoV infections in human and mouse lung cell cultures, we found three MERS-CoV datasets in GEO for use in our study (Table 2). GSE81909 and GSE100504 examined cultures of primary human airway epithelial cells infected with mock or wild type MERS-CoV (icMERS). Gene expression data for these datasets were profiled on the Probe Name version of the commercial Agilent-026652 Whole Human Genome Microarray 4×44K v2 (GEO GPL13497). GSE108594 had mouse lung cell cultures that were mock or MERS-CoV infected across an inoculation dose range (0, 10^4^, 10^5^, and 10^6^) and profiled on the Probe Name version of Agilent-026655 Whole Mouse Genome Microarray 4×44K v2 (GEO GPL11202). For all MERS-CoV datasets, expression data provided by GEO were already normalized so we selected data for the 48hr post-infection time point as needed and cleaned data as previously described for SARS-CoV datasets.

**Table 2.** MERS-CoV and SARS-CoV2 Infection Datasets Utilized for this Study

| Dataset | Description | Platform | Probes | IDs | Genes |
| --- | --- | --- | --- | --- | --- |
| GSE81909<sup>1</sup> | Log2 normalized cultures of primary human airway epithelial cells infected with mock or 5 plaque-forming units/cell of wild type MERS-CoV (icMERS) | GPL13497 | 34184 | 29833 | 26049 |
| GSE100504<sup>1</sup> | Log2 normalized cultures of primary human airway epithelial cells infected with mock or 5 plaque-forming units/cell of wild type MERS-CoV (icMERS) | GPL13497 | 34184 | 29833 | 26049 |
| GSE108594<sup>1</sup> | Quantile normalized cultures of mouse lung cells infected with mock, 10<sup>4</sup>, 10<sup>5</sup>, or 10<sup>6</sup> doses of 3 plaque-forming units wild-type MERS-CoV | GPL11202 | 39485 | 33244 | 31895 |
| GSE152586 | Unnormalized<sup>2</sup> samples of human alveolar type II cell lung organoids 48hrs post mock or SARS-CoV2 infection | GPL20795<br>GPL28694 | 26496 | 26496 | 26494 |
| GSE160435 | Unnormalized<sup>2</sup> human lung organoid samples generated from primary lung alveolar type II cells 48hrs post mock or SARS-CoV2 infection | GPL24676<br>GPL29320 | 60666 | 60666 | 60598 |
| GSE155518 | Unnormalized<sup>2</sup> human lung organoid samples generated from primary lung alveolar type II cells 48hrs post mock or SARS-CoV2 infection | GPL24676 | 60765 | 60765 | 60686 |

<sup>1</sup>Datasets collected samples over a time course. 48hr samples were used.
<sup>2</sup>Samples were z-scored across all dataset samples regardless of time or infection for normalization before use.
Platform, Gene Expression Omnibus platform identifiers. Probes, number of platform probes with expression data in dataset. IDs, number of probes with Gene Expression Omnibus provided Entrez identifiers. Genes, number of unique Entrez identifiers used for gene signature definition with representatives for duplicate Entrez identifiers being selected by highest coefficient of variation. Values provided in the Genes column reflect the number of genes in gene signatures defined for this study.

#### 2.1.3 SARS-CoV2 Datasets

To compare molecular changes identified in SARS-CoV to changes associated with SARS-CoV2 infections in human lung cell cultures, we found three SARS-CoV2 datasets in GEO with mock or SARS-CoV2 infected samples collected at 48hrs for use in our study (Table 2). GSE152586 contained gene expression data for cultures of human alveolar type II cell organoids [55] that were profiled on HiSeq X Ten (GEO GPL20795 and GPL28694). GSE160435 and GSE155518 contained gene expression data from organoids generated from human primary alveolar type II cells and were profiled on Illumina NovaSeq 6000 (GEO GPL24676 and/or GPL29320). GEO had no available SARS-CoV2 datasets utilizing mouse cultures or samples appropriate for use in this study. All provided SARS-CoV2 expression data was z-score normalized and cleaned as previously described for SARS-CoV datasets.

### 2.2 Defining Gene Signatures

We measured differential gene expression by Welch’s two-sample T-test score of normalized values to generate gene signatures (*i.e.*, gene lists ranked by differential gene expression between SARS and mock infected samples). Genes that were over-expressed in SARS- compared to mock infected samples (*e.g.*, positive T-score) fall within the positive tail of the gene signature while under-expressed genes (*e.g.*, negative T-score) fall in the negative tail (Figure 1A). Genes with no substantial change in expression between SARS and mock infected samples (*e.g.*, T-score around 0) were located toward the middle of the gene signature. Therefore, genes that fall within the tails of a gene signature likely changed in response to a specific SARS infection. We noted that three signatures were substantially skewed (*i.e.*, rank of genes where T-score crosses from positive to negative was in top or bottom quartile of signature). For substantially skewed signatures, we adjusted all T-scores in the signature by the T-score of the gene at mid-point so that genes in the signature were balanced between positive and negative T-scores.

**Figure 1.**
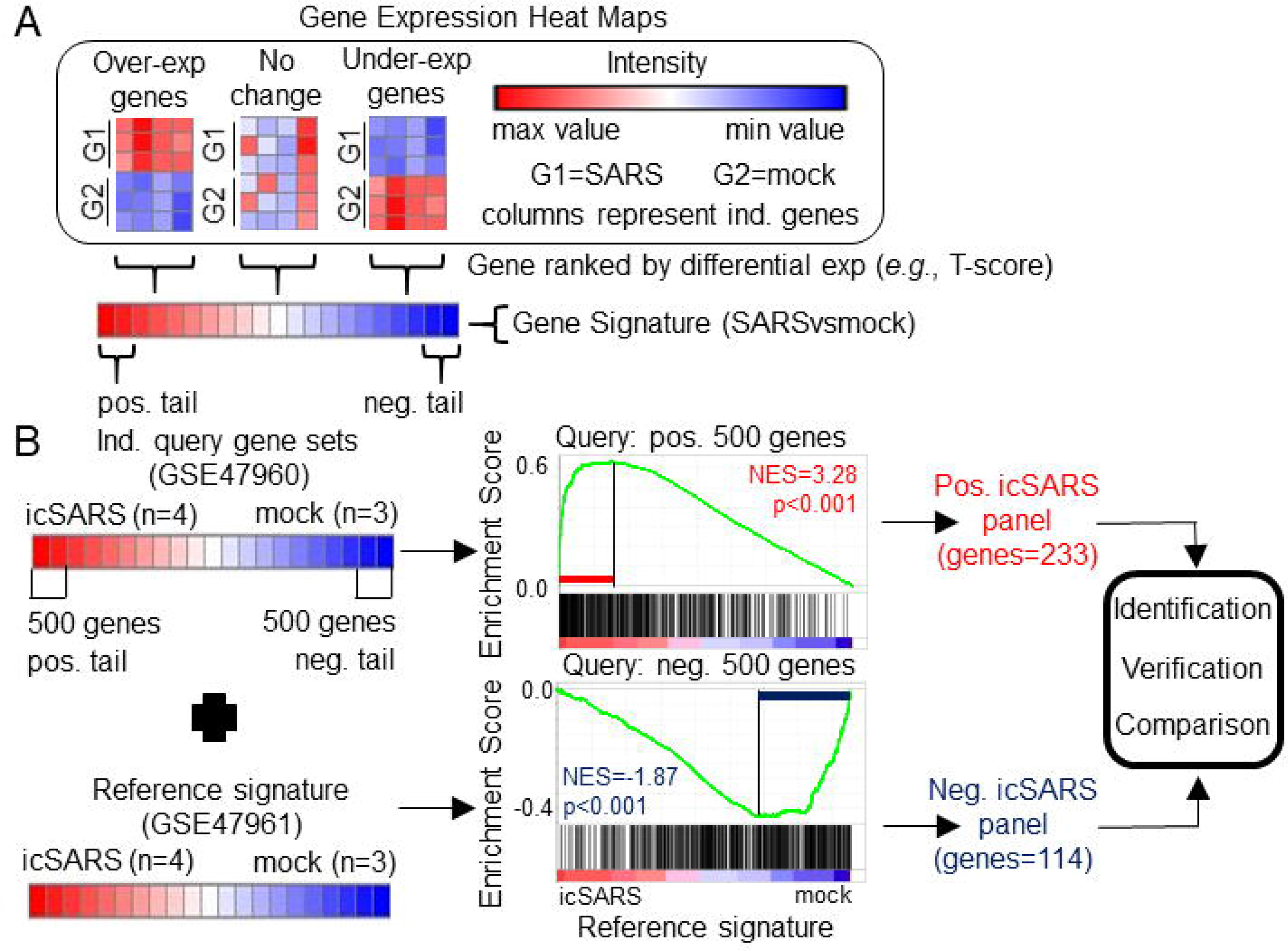
Gene Signature Definition and Generation icSARS Gene Panels A) Schematic definition of a gene signature. Differences in gene expression between two groups, such as SARS and mock infected lung cells, are measured by Welch’s two-sample TLtest score. Gene signatures are ranked lists of genes from high (red) to low (blue) differential mRNA expression between groups. B) Generation of icSARS gene panels for use in this study. To identify differentially expressed genes associated with icSARS infection in human airway epithelial cell cultures, query gene sets containing either the 500 most over- or under-expressed genes from positive or negative tails of the gene signature generated from the Gene Expression Omnibus (GEO) accession number GSE47960 mRNA expression dataset. The positive and negative tail query sets were compared individually to the gene signature generated from the GEO GSE47961 dataset, which was used as reference for Gene Set Enrichment Analysis (GSEA). From this GSEA computed two enrichment plots, one for each query set, and their associated normalized enrichment score (NES) and p-value which represent the extent of enrichment between query set and reference signature. GSEA also identified leading-edge genes, which are genes that contribute most to achieving maximum enrichment. Two gene panels were defined from leading-edge genes identified in each query set. These gene panels were used in this study for three purposes: 1) identification of gene expression changes associated with icSARS infection in human airway epithelial cell cultures, 2) verification of identified findings in independent datasets, and 3) comparison to other gene signatures representing changes in gene expression associated with other SARS infections.

From the 17 SARS datasets previously described, we defined a total of 37 gene signatures. We generated 29 SARS-CoV gene signatures, which included 17 gene signatures from human lung cultures and 12 signatures from mouse lung samples (Table 3). Specifically, we defined seven icSARSvsmock signatures (six in human cultures, one in mouse samples), one Urbanivsmock signature with the Urbani strain in human cultures, eight MA15vsmock signatures doses in mouse samples representing varying inoculation, five ORF6vsmock signatures (four in human cultures, one in mouse samples), five BATSRBDvsmock signatures (four in human cultures, one in mouse samples), two NSP16vsmock signatures (one in human cultures, one in mouse samples), and one ExoNIvsmock signature from human cultures. For MERS-CoV datasets, we defined five gene signatures that included two gene signatures from human lung cultures and three signatures from mouse lung samples (Table 4). For SARS-CoV2 datasets, we defined three gene signatures from human lung cultures (Table 4).

**Table 3.** SARS-CoV Gene Signatures Defined in this Study

| Dataset | Group 1 (N) | Group 2 (N)<sup>1</sup> | Gene signature | Use | High<sup>2</sup> | Low<sup>2</sup> | Cross<sup>3</sup> |
| --- | --- | --- | --- | --- | --- | --- | --- |
| GSE47960 | icSARS (4) | mock (3) | icSARSvsmock | I | 12.5 | -13.2 | 7245 |
|  | dORF6 (4) | mock (3) | ORF6vsmock | C | 12.5 | -13.2 | 8738 |
|  | BatSRBD (3) | mock (3) | BATSRBDvsmock | C | 11.6 | -12.1 | 8076 |
| GSE47961 | icSARS (4) | mock (3) | icSARSvsmock | I | 12.0 | -13.1 | 8285 |
|  | dORF6 (4) | mock (3) | ORF6vsmock | C | 13.1 | -12.1 | 9026 |
|  | BatSRBD (4) | mock (3) | BATSRBDvsmock | C | 12.1 | -6.7 | 8421 |
| GSE47962 | icSARS (3) | mock (3) | icSARSvsmock | V | 12.1 | -12.1 | 8703 |
|  | dORF6 (3) | mock (3) | ORF6vsmock | C | 12.2 | -12.2 | 8225 |
|  | BatSRBD (3) | mock (3) | BATSRBDvsmock | C | 12.2 | -12.2 | 9157 |
| GSE37827 | icSARS (3) | mock (3) | icSARSvsmock | V | 12.2 | -12.2 | 8261 |
|  | BatSRBD (3) | mock (3) | BATSRBDvsmock | C | 12.2 | -11.8 | 8843 |
| GSE48142 | icSARS (3) | mock (3) | icSARSvsmock | V | 12.2 | -12.2 | 6918 |
|  | ExoNI (3) | mock (3) | ExoNIvsmock | C | 12.2 | -9.9 | 7570 |
|  | dNSP16 (3) | mock (3) | NSP16vsmock | C | 12.2 | -12.2 | 6638 |
| GSE33267 | icSARS (3) | mock (3) | icSARSvsmock<sup>4</sup> | V | 15.9 | -8.6 | 9874 |
|  | dORF6 (3) | mock (3) | ORF6vsmock | C | 12.2 | -12.2 | 7378 |
| GSE17400 | Urbani (3) | mock (3) | Urbanivsmock | C | 12.2 | -12.2 | 10571 |
| GSE50000 | icSARS (5) | mock (4) | icSARSvsmock | C | 6.8 | -7.7 | 10007 |
|  | MA10<sup>5</sup> (4) | mock (4) | MA10<sup>5</sup>vsmock | C | 9.4 | -10.2 | 9882 |
|  | MA10<sup>4</sup> (4) | mock (4) | MA10<sup>4</sup>vsmock | C | 14.1 | -14.1 | 9759 |
|  | BatSRBD (5) | mock (4) | BATSRBDvsmock | C | 9.9 | -9.5 | 10185 |
| GSE33266 | MA10<sup>5</sup> (5) | mock (3) | MA10<sup>5</sup>vsmock<sup>4</sup> | C | 13.6 | -14.6 | 10803 |
|  | MA10<sup>4</sup> (5) | mock (3) | MA10<sup>4</sup>vsmock | C | 14.0 | -14.1 | 11654 |
|  | MA10<sup>3</sup> (5) | mock (3) | MA10<sup>3</sup>vsmock<sup>4</sup> | C | 12.8 | -6.6 | 10803 |
|  | MA10<sup>2</sup> (5) | mock (3) | MA10<sup>2</sup>vsmock<sup>4</sup> | C | 7.3 | -6.4 | 10803 |
| GSE49262 | MA10<sup>5</sup> (3) | mock (3) | MA10<sup>5</sup>vsmock | C | 4.5 | -2.5 | 10073 |
|  | dORF6 (3) | mock (3) | ORF6vsmock | C | 4.4 | -2.5 | 10211 |
| GSE49263 | MA10<sup>5</sup> (4) | mock (3) | MA10<sup>5</sup>vsmock | C | 4.8 | -2.5 | 10384 |
|  | dNSP16 (4) | mock (3) | NSP16vsmock | C | 5.0 | -2.6 | 10188 |

<sup>1</sup>Same mock samples used across signatures in same dataset.
<sup>2</sup>These values define the range of Welch's two-sample T-scores that were used to rank genes in the signature.
<sup>3</sup>Values in the Cross column reflect where (*i.e.*, which gene) in the signature the T-score crosses 0.
<sup>4</sup>T-scores in gene signature adjusted so Cross (T-score=0) occurs near center of the signature.
C, comparison; I, identification; N, number of samples; V, verification.

**Table 4.** MERS-CoV and SARS-CoV2 Gene Signatures Defined in this Study

| Dataset | Group 1 (N) | Group 2 (N)<sup>1</sup> | Gene signature | Use | High<sup>2</sup> | Low<sup>2</sup> | Cross<sup>3</sup> |
| --- | --- | --- | --- | --- | --- | --- | --- |
| GSE81909 | icMERS (5) | mock (5) | icMERSvsmock | C | 5.1 | -1.7 | 11451 |
| GSE100504 | icMERS (5) | mock (5) | icMERSvsmock | C | 5.8 | -2.5 | 12836 |
| GSE108594 | MERS (4) | mock (4) | MERS10<sup>4</sup>vsmock | C | 4.3 | -1.5 | 14276 |
|  | MERS (4) | mock (4) | MERS10<sup>5</sup>vsmock | C | 4.2 | -1.8 | 14593 |
|  | MERS (4) | mock (4) | MERS10<sup>6</sup>vsmock | C | 4.1 | -2.0 | 15545 |
| GSE152586 | SARS2 (3) | mock (3) | SARS2vsmock | C | 12.2 | -12.2 | 13084 |
| GSE160435 | SARS2 (5) | mock (5) | SARS2vsmock | C | 15.8 | -12.1 | 26969 |
| GSE155518 | SARS2 (3) | mock (3) | SARS2vsmock | C | 13.7 | -10.8 | 30343 |

<sup>1</sup>Same mock samples used across signatures in same dataset.
<sup>2</sup>These values define the range of Welch's two-sample T-scores that were used to rank genes in the signature.
<sup>3</sup>Values in the Cross column reflect where (*i.e.*, which gene) in the signature the T-score crosses 0.
<sup>4</sup>T-scores in gene signature adjusted so Cross (T-score=0) occurs near center of the signature.
C, comparison; N, number of samples

### 2.3 Overview of Gene Set Enrichment Analysis

We used Gene Set Enrichment Analysis (GSEA) as a statistical method that estimated enrichment between a query gene set (*i.e.*, unranked list of genes) and a reference gene signature [40]. GSEA used T-score to calculate a running summation enrichment score where hits (*i.e.*, matches between query set and reference signature) increased the enrichment score proportional to the ranking statistical metric (*e.g.*, T-score) and a miss (*i.e.*, non-matches between query set and reference signature) decreased the enrichment score. From this, GSEA determined a maximum enrichment score for the specific query set and reference signature. Leading-edge genes contributed to reaching the maximum enrichment score, indicating leading-edge genes were associated with cellular response to a specific β-coronavirus infection. Further, GSEA calculated a normalized enrichment score (NES) from 1000 permutations of the reference signature to estimate the significance of enrichment between a specific query set and reference signature. This work used the javaGSEA Desktop Application release 3.0 version of GSEA available from Broad Institute to perform gene T-ranking for signature formation and GSEA for gene identification, verification, and comparison.

### 2.4 Identification of icSARS Associated Genes

To identify gene expression changes associated with icSARS infection, we generated two icSARS gene panels (Figure 1B). To do this, we selected 500 genes from the positive and negative tails from the GSE47960-derived icSARSvsmock gene signature and used them to form two individual query gene sets. GSEA compared each query gene set to the GSE47961-derived icSARSvsmock gene signature (reference). Leading-edge genes from each analysis were used to define the two icSARS gene panels, one panel per tail, and we identified genes included in icSARS panels as being associated with icSARS infection. Pathway enrichment analysis was performed on both icSARS gene panels using Database for Annotation, Visualization and Integrated Discovery (DAVID) v6.8. DAVID was a web-accessible knowledgebase with a comprehensive set of functional annotation tools for researchers to understand biological meaning behind large list of genes [56, 57].

### 2.5 Verification of icSARS Gene Panels

To verify the icSARS gene panels, we performed GSEA between icSARS gene panels and GSE47962-derived, GSE37827-derived, GSE48142-derived, and GSE33267-derived icSARSvsmock signatures. To assess if results generated from GSEA could be achieved randomly, we randomly selected 1000 gene panels consisting of either 233- or 114-genes, to match the number of genes in the positive and negative icSARS panels, respectively, from the GPL6480 platform. These analyses generated a null distribution of NES to which we compared the NES achieved by icSARS gene panels for each reference gene signature and counted the number of equal or better NES to estimate significance (*i.e.*, distribution p-value). Histogram data and associated graphs (*e.g.*, distribution curves and box and whiskers plot) were generated using XLStat version 2020.3 [58, 59], which was a user-friendly, commercial, data analysis add-on for Microsoft Excel.

### 2.6 Comparison to icSARS-induced Gene Expression Changes in Mice and Across Other SARS Strains

To examine differential gene expression of genes from the icSARS gene panels in mice, we performed GSEA between icSARS gene panels (queries) and the GSE50000-derived icSARSvsmock signature (reference). We compared gene membership across leading-edges identified in the analysis of icSARS verification signatures and the icSARS mouse signature to identify genes associated with icSARS infection between models. Any icSARS panel genes that were not included the dataset’s platform still received consideration when examining across identified leading-edge genes. To expand this analysis by comparing gene signatures across infections caused by other SARS-CoV strains, we repeated this GSEA-based meta-analysis approach on infections with other strains, such as Urbani, MA15, SARS-BatSRBD, and mutants in ORF6, NSP16, or ExoNI. All SARS-CoV strains except ExoNI had strain-matched samples in both mouse samples and human lung cultures. We included MA15-infected mouse signatures over a range of inoculation doses (Table 3) to mimic the range of infection severity that would be encountered clinically (*i.e.*, asymptomatic to severe patient presentation). Genes associated with SARS-CoV infection were identified by gene membership across leading-edges identified in these 22 analyses. Further, we repeated this GSEA-based meta-analysis approach to include gene signatures derived from MERS-CoV and SARS-CoV2 infections to compare gene signatures across major SARS infectious strains and examine gene membership across leading-edges. Random modelling for all comparisons were done as previously described. Heat maps were generated by a user-friendly, web-based program from Broad Institute that produces customizable heat maps named Morpheus, https://software.broadinstitute.org/morpheus.

## 3 Results

### 3.1 Gene Signature Approach Identified Gene Expression Changes Associated with icSARS Infection for Human Lung Epithelium Cells *in vitro*

To identify genes associated with response to an icSARS infection, we first defined the GSE47960-derived and GSE47961-derived icSARSvsmock gene signatures (Table 2). We used the GSE47960-derived icSARSvsmock to generate two gene sets containing the 500 most differentially expressed genes from the positive and negative tails of GSE47960-derived icSARSvsmock (T-score >2.9 and <−3.2 for positive and negative tails, respectively). We chose 500 genes to capture maximum coverage of the signature that was allowable by GSEA [40]. To assess similarity between these two gene signatures, we calculated enrichment using GSEA between either GSE47960-derived icSARSvsmock positive or negative tail gene sets and the GSE47961-derived icSARSvsmock and achieved NES=3.28 and NES=-1.87 for positive and negative tail query gene sets, respectively, both with a GSEA p-value<0.001. We defined separate positive and negative icSARS gene panels from the 233 and 114 leading-edge genes identified (Supplemental Material STable 1), representing over- and under-expressed genes associated with icSARS infection.

To explore potential factors that could impact our results, we wanted to see if similar results were generated from panels defined using 1) reversed query and reference signatures, 2) different query set sizes, and 3) different time points. First, we repeated the process of defining panels except reversing the query and reference signatures by using the 500-gene tails from GSE47961-derived icSARSvsmock signature and GSE47960-derived icSARSvsmock, respectively. We noted that the size of the positive panel did not substantially change from 233 genes (Table 1) to 240 genes (Supplemental Material STable 2). However, we noted that the size of the negative icSARS panel did change substantially from 114 to 331 genes. Despite this substantial change in size, we observed that enrichment did not substantially change for either positive or negative tail query gene sets when reversed (Supplemental Material SFig 1). From this we conclude that reversing which signature was used as reference and to derive query gene sets does not change the overall result. Next, we repeated the process of defining panels except we used smaller query sizes (100, 200, 300, and 400 genes). We found similar enrichment regardless of query size (Supplemental Material SFig 1), highlighting the strong similarity between these signatures. Finally, we examined how time point selection (24, 48, 72, and 96hrs) may impact our results. We found reproducible, statistically significant enrichment beginning at 48hrs, suggesting that 48hrs was the earliest timepoint where gene expression changes could be reliably detected. Based on these results collectively, we focused our meta-analysis on the 48hr positive (233 genes) and negative (114 genes) icSARS gene panels defined from using the 500-gene tails of GSE47960-derived icSARSvsmock gene signature as query, since identified leading-edge genes represented over- and under-expressed genes associated with icSARS infection at the earliest time point with consistent detectable gene signature similarities.

Among genes in positive icSARS panel, we found 13 genes (5.5% of the panel) had previously reported associations with icSARS infections via single-gene analysis with minimum fold change of 2.0 and maximum false discovery rate-corrected p-value of 0.05 between mock and icSARS time-matched samples [30], demonstrating the predictive ability of our gene signature approach. There were no genes in the negative icSARS gene panel with established associations with icSARS infection. We also identified genes with no prior association with icSARS infection. These genes included 12 genes had a zinc finger in addition to the two that were already reported (5.2% of the panel), six genes encoding IFN-induced proteins (2.6%), five genes from the solute carrier family (2.1%), and six genes encoding for uncharacterized proteins (2.6%) in the positive icSARS gene panel. In the negative icSARS gene panel, we found three zinc finger protein genes (2.6%), three genes from the solute carrier family (2.6%), and five genes encoding for uncharacterized proteins (4.4%). We speculated that these identified genes without previously reported associations with icSARS infections in human lung epithelium cultures were also associated with an icSARS infection.

To expand on our analysis, we examined the cellular roles that genes in the icSARS panels were involved in. We used DAVID to calculate enrichment between the icSARS panels and pathways in commonly used knowledgebases. We noted that GO Biological Processes (BP) database returned the most identified significantly enriched pathways compared to other databases (data not shown), so we focused this discussion on GO-BP data to avoid confusion from overlapping pathway and gene inclusion variations across different knowledgebases. DAVID identified 49 significant GO-BP pathways (EASE score p-value<0.05) from the positive icSARS panel and 10 significant pathways from the negative icSARS panel (Supplemental Material STable 3). Most significantly enriched pathways have experimentally established associations with icSARS and other β-coronavirus infections, such as up-regulation of type I IFN signaling pathway (GO:0060337, p-value<0.001), NF-kappaB processes (transcription factor activity: GO:0051092, p-value=0.001; signaling: GO:0043123, p-value=0.043), p38MAPK cascade (GO:1900745, p-value=0.015) and apoptotic process (GO:0006915, p-value=0.018) and down-regulation of cilium morphogenesis (GO: 0060271, p-value=0.038) and assembly (GO: 0042384, p-value=0.0.030), demonstrating our gene signature approach’s ability to detect pathways associated with response to an icSARS infections [60–62]. We also find several pathways, such as SMAD protein signal transduction (GO:0060395, p-value=0.037), with no prior associations to icSARS infection though they have been identified in mouse infections with the dORF strain [2]. Therefore, we speculated that pathways without prior association to icSARS infections identified here were also involved in response to an icSARS infection.

### 3.2 Enrichment of icSARS Gene Panels and Specific icSARS Panel Genes Verified in Independent Datasets

To verify our icSARS gene panels were associated with response to an icSARS infection, we used GSEA to calculate enrichment between our icSARS panels and four icSARSvsmock verification gene signatures derived from independent datasets (Table 2). We found significant similarity between positive icSARS panels and all icSARSvsmock verification signatures (Figures 2A through 2D, NES>2.95, GSEA p-value <0.001). We also observed significant similarity between negative icSARS panels and the icSARSvsmock verification signatures (Figures 2E through 2H, NES<−2.14, p-value <0.001). To determine how likely the NES achieved for icSARS panels would be achieved by random chance, we generated a random model to match the size and potential composition of our icSARS panels. We observed NES achieved by our icSARS panels were consistently outside the range of NES generated randomly (null distribution p-value<0.001, Figures 2I through 2L), illustrating that NES achieved by our icSARS panels were non-random. Taken together, these results demonstrated that the enrichment achieved from our icSARS panels was true and reproducible.

**Figure 2.**
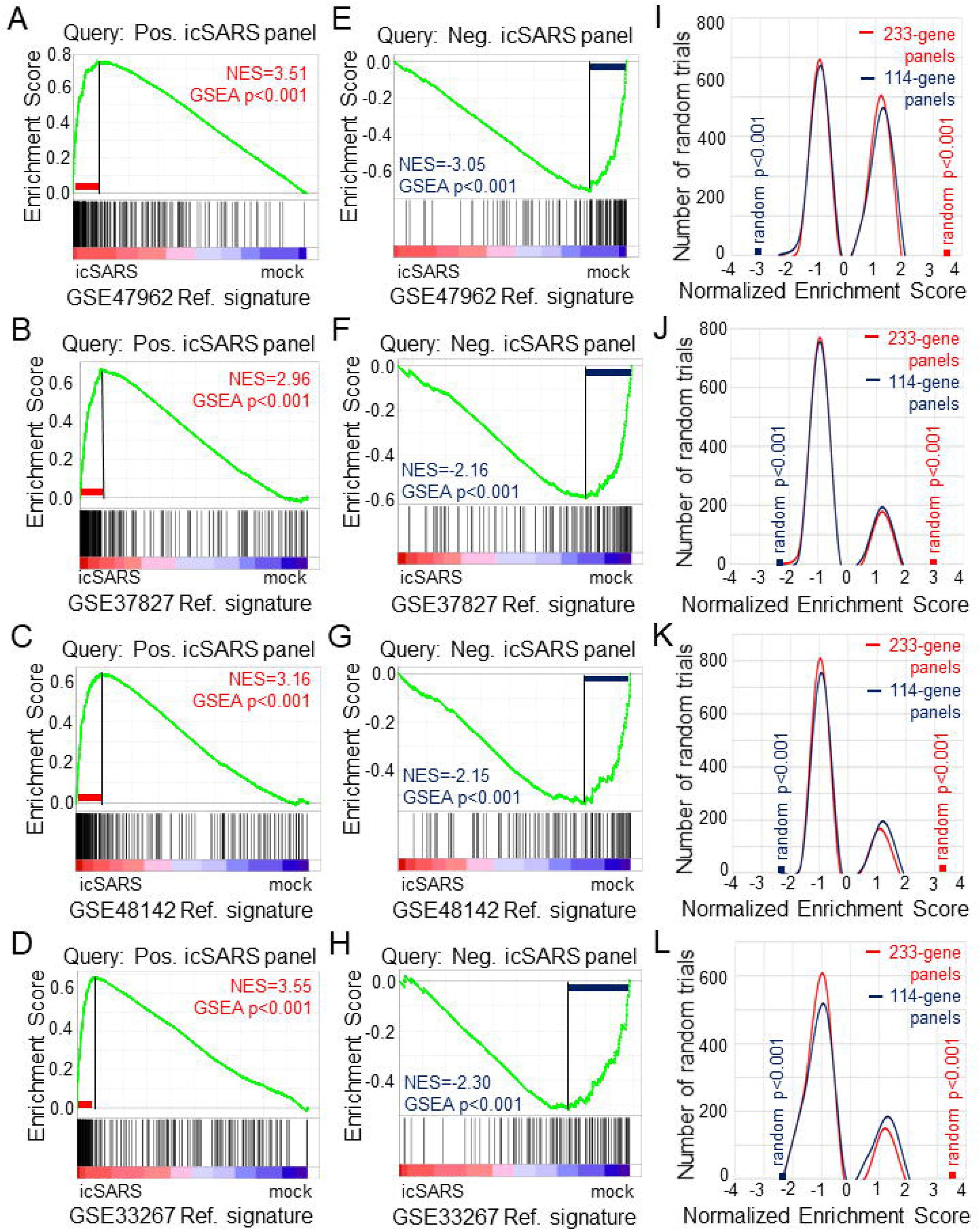
Verification of icSARS Gene Panels in Independent Datasets A) Gene Set Enrichment Analysis (GSEA) calculated enrichment, as determined by normalized enrichment score (NES), between the positive icSARS gene panel and the GSE47962-derived icSARSvsmock gene signature. B) GSEA between the positive icSARS gene panel and GSE37827-derived icSARSvsmock gene signature. C) GSEA between the positive icSARS gene panel and GSE48142-derived icSARSvsmock gene signature. D) GSEA between the positive icSARS gene panel and GSE33267-derived icSARSvsmock gene signature. E) GSEA between the negative icSARS panel and the GSE47962-derived icSARSvsmock signature. F) GSEA between the negative icSARS panel and the GSE37827-derived icSARSvsmock signature. G) GSEA between the negative icSARS panel and the GSE48142-derived icSARSvsmock signature. H) GSEA between the negative icSARS panel and the GSE33267-derived icSARSvsmock signature. I) Distribution plot of NES from 1000 randomly generated gene panels (individual queries) compared to the GSE47962-derived icSARSvsmock signature. J) Distribution plot of NES from 1000 randomly generated gene panels compared to the GSE37827-derived icSARSvsmock signature. K) Distribution plot of NES from 1000 randomly generated gene panels compared to the GSE48142-derived icSARSvsmock signature. L) Distribution plot of NES from 1000 randomly generated gene panels compared to the GSE33267-derived icSARSvsmock signature.

To determine which icSARS panel genes were verified across all signatures, we examined leading-edge genes identified by GSEA for each verification signature (Supplemental Material STable 4). We observed 51 genes from the positive icSARS panel and 22 genes from the negative icSARS panel were shared across all verification signatures. Of the 51 positive icSARS panel genes, we noted 10 genes had reported associations with icSARS infection via single-gene analysis with minimum fold change of 2.0 and maximum false discovery rate-corrected p-value of 0.05 between mock and icSARS time-matched samples [30]: nuclear factor of kappa light polypeptide gene enhancer in B-cells inhibitor, alpha (Entrez ID 4792), zinc finger CCCH-type containing 12A (ID 80149), cysteine-serine-rich nuclear protein 1 (ID 64651), zinc finger protein 433 (ID 163059), hairy and enhancer of split 1 from Drosophila (ID 3280), transformer 2 beta homolog (ID 6434), V-rel reticuloendotheliosis viral oncogene homolog B (ID 5971), pim-3 oncogene (ID 415116), chemokine (C-X-C motif) ligand 2 (ID 2920), and C-X-C motif chemokine ligand 10 (ID 3627). These data together supported the conclusion that our shared leading-edge genes were associated with icSARS infection in human lung cultures and supported the hypothesis that identified genes without previously reported associations were also associated with icSARS infection in human lung cultures.

### 3.3 Positive icSARS Gene Panel Enriched in Mouse-derived icSARS Gene Signature

To determine if icSARS gene panels were also associated with response to an icSARS infection in a mouse model, we used GSEA to calculate enrichment between icSARS panels and the GSE50000-derived icSARSvsmock gene signature (Table 3). We observed significant enrichment with the positive icSARS panel (NES=1.54, GSEA p-value<0.001, Figure 3A) but not the negative icSARS panel (NES=0.81, p-value=0.848, Figure 3B). Achieved NES were non-random for the positive icSARS panel (NES range: −1.65 to 1.57, null distribution p-value=0.002, Figure 3C), but not for the negative icSARS panel (NES range: −1.72 to 1.64, p-value=0.601). These results suggested that genes in the positive icSARS panel were associated with an icSARS infection in both in human lung cultures and mouse lung samples, but the same cannot be concluded for the negative icSARS panel. Therefore, we did not consider leading-edge genes identified by GSEA from the negative icSARS panel.

**Figure 3.**
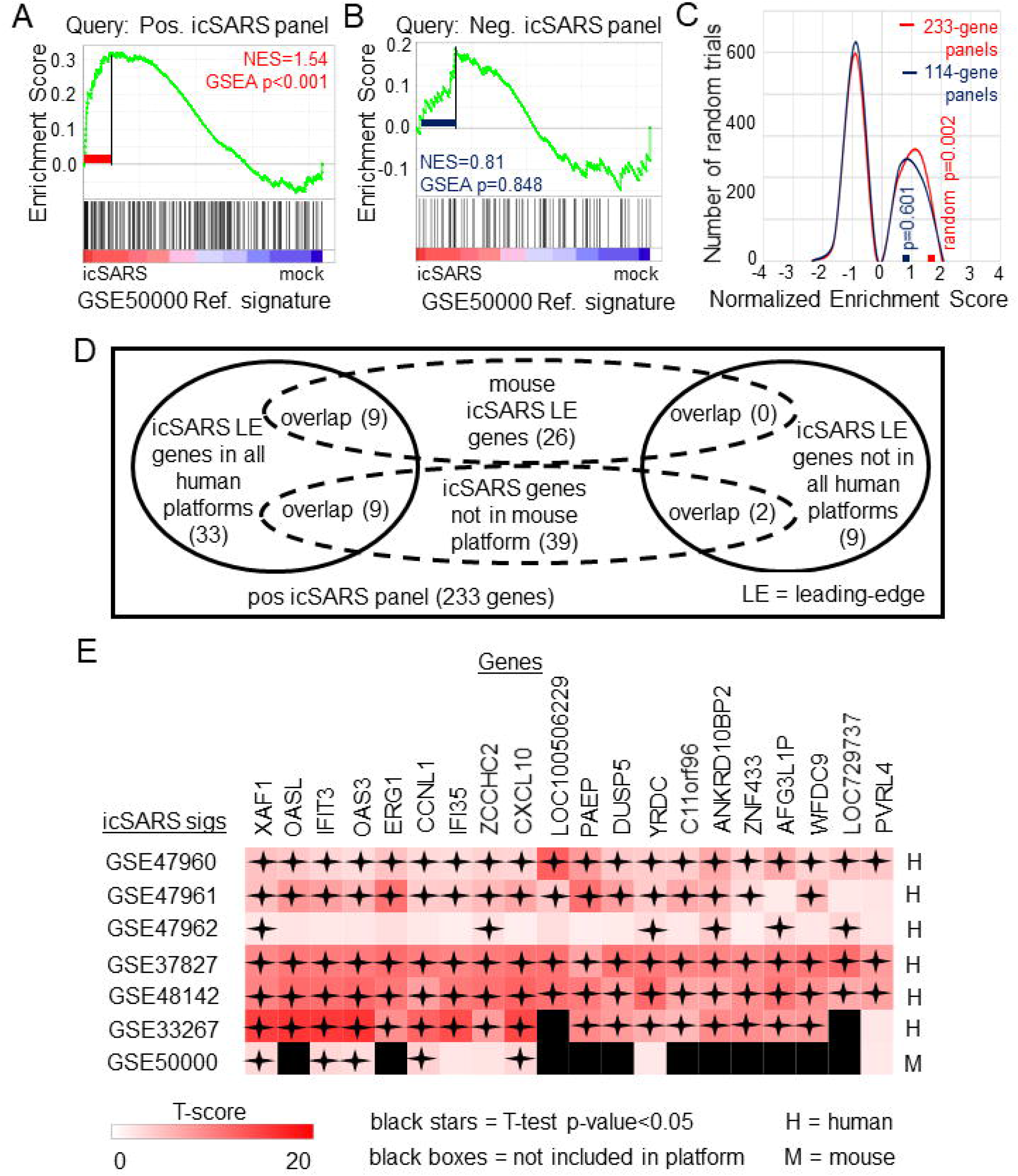
Positive icSARS Panel Enrichment in icSARS Infected Mouse Model Revealed Genes Associated with icSARS Infection A) Gene Set Enrichment Analysis (GSEA) calculated enrichment, as determined by normalized enrichment score (NES), between the positive icSARS gene panel and the GSE50000-derived icSARSvsmock gene signature. B) GSEA between the negative icSARS panel and the GSE50000-derived icSARSvsmock signature. C) Distribution plot of NES from 1000 randomly generated gene panels (individual queries) compared to the GSE50000-derived icSARSvsmock signature (reference). D) Venn diagram of the inclusion and overlap of positive icSARS panel genes in identified leading-edges and dataset platforms across icSARS-CoV human and mouse gene signatures. E) Heat map of T-scores for the 20 positive icSARS panel leading-edge genes identified in (D).

When examining across leading-edge genes identified by GSEA from the positive icSARS panel (Supplemental Material STable 5), our meta-analysis approach found nine of the 51 genes verified across human lung cultures were also in the leading-edge from mouse lung samples (Figure 3D): IFN-induced protein with tetratricopeptide repeats 3 (human Entrez ID: 3437, mouse Entrez ID: 15959, gene symbol: IFIT3), C-X-C motif chemokine ligand 10 (human: 3627, mouse: 15945, symbol: CXCL10), XIAP associated factor 1 (human: 54739, mouse: 327959, symbol: XAF1), 2′-5′-oligoadenylate synthetase 3 100kDa (human: 4940, mouse: 246727, symbol: OAS3), zinc finger CCHC domain containing 2 (human: 54877, mouse: 227449, symbol: ZCCHC2), IFN-induced protein 35 (human: 3430, mouse: 70110, symbol: IFI35), yrdC domain containing (human: 79693, mouse: 230734, symbol: YRDC), poliovirus receptor-related 4 (human: 81607, mouse: 71740, symbol: PVRL4), and cyclin L1 (human: 57018, mouse: 56706, symbol: CCNL1). We also noted that several icSARS panel genes were not included in the mouse dataset’s platform (Supplemental Material STable 6) with nine of these genes verified across leading-edges for icSARS gene signatures derived from human cell cultures and two genes not included in all verification icSARS signatures. We did not exclude genes from further consideration based on platform inclusion. A heat map of differential gene expression (*i.e.*, T-scores) for these 20 genes across all six icSARS gene signatures revealed that only XAF1 was statistically significant (Welch’s two-sampled, two-sided T-test p-value<0.05) in all icSARS gene signatures (Figure 3E), highlighting the advantage of using a GSEA-based rather than single-gene (*e.g.*, T-score only) meta-analysis, which would have likely missed the other genes identified here because of borderline significance in at least one signature.

### 3.4 Positive icSARS Gene Panel Significantly Enriched Across Other SARS-CoV Strains

After identifying and verifying genes associated with an icSARS infection, we wanted to find differential gene expression similarities between icSARS infection and infections from other SARS-CoV stains with varying levels of virulence [30, 32–35]. Identified gene similarities represented genes associated with a SARS-CoV infection regardless of virulence, and we hypothesized that shared genes may be useful targets clinically to preclude or overcome SARS infection. To identify genes associated with SARS-CoV infection across strains, we used GSEA to compare icSARS gene panels to 18 gene signatures derived from samples of human lung cultures or mouse lung samples mock or SARS-CoV infected with one of six strains (Urbani, MA15, dORF6, BAT-SRBD, dNSP16, and ExoNI; Table 2). Urbani and MA15 were fully virulent, while BAT, dORF6, dNSP and ExoNI were attenuated by established differences in host range or gene mutation mechanisms. These strains were selected so all available GEO datasets containing SARS-CoV and mock samples collected at 48hrs post infection were included in our meta-analysis. We found the positive icSARS panel significantly enriched (NES>1.66, GSEA p-value<0.01) across all 18 gene signatures (Figure 4A). We confirmed via random modelling as previously described for all icSARS signatures that achieved significant positive NES were non-random (null distribution p-value<0.002, Figure 4B), except for the GSE33266-derived MA15(10^2^)vsmock signature which was the lowest infectious dose examined in this study. Due to this finding, we remove the MA15(10^2^)vsmock signature from further analysis. The negative icSARS panel was significantly enriched across most signatures (NES<-1.52, p-value<0.01) except two BATvsmock signatures, the GSE47961-derived signature from human cultures (NES=-0.95, p-value=0.569) and the GSE50000-derived signature from mouse samples (NES=-0.81, p-value=0.846), and the GSE33266 MA15(10^3^)vsmock signature from mouse samples (NES=0.90, p-value=0.635). Random modelling with gene sets the same size as the negative icSARS panel supported enrichment findings observed with the negative icSARS panel since enriched signatures were not randomly enriched (Figure 4C, p-value<0.002) but unenriched signatures were randomly enriched (p-value>0.075). As additional verification of our findings, we re-ran GSEA using the panels generated from GSE47961-derived icSARSvsmock (500-gene queries), and noticed this subtle change did not affect observed enrichment across 27 signatures (Supplemental Material SFig 2) Taken together, these data supported our previous findings of consistent enrichment of the positive icSARS panel and inconsistent enrichment of the negative icSARS panel across icSARS signatures, suggesting that genes shared across positive icSARS panel leading-edges were associated with SARS-CoV infections generally.

**Figure 4.**
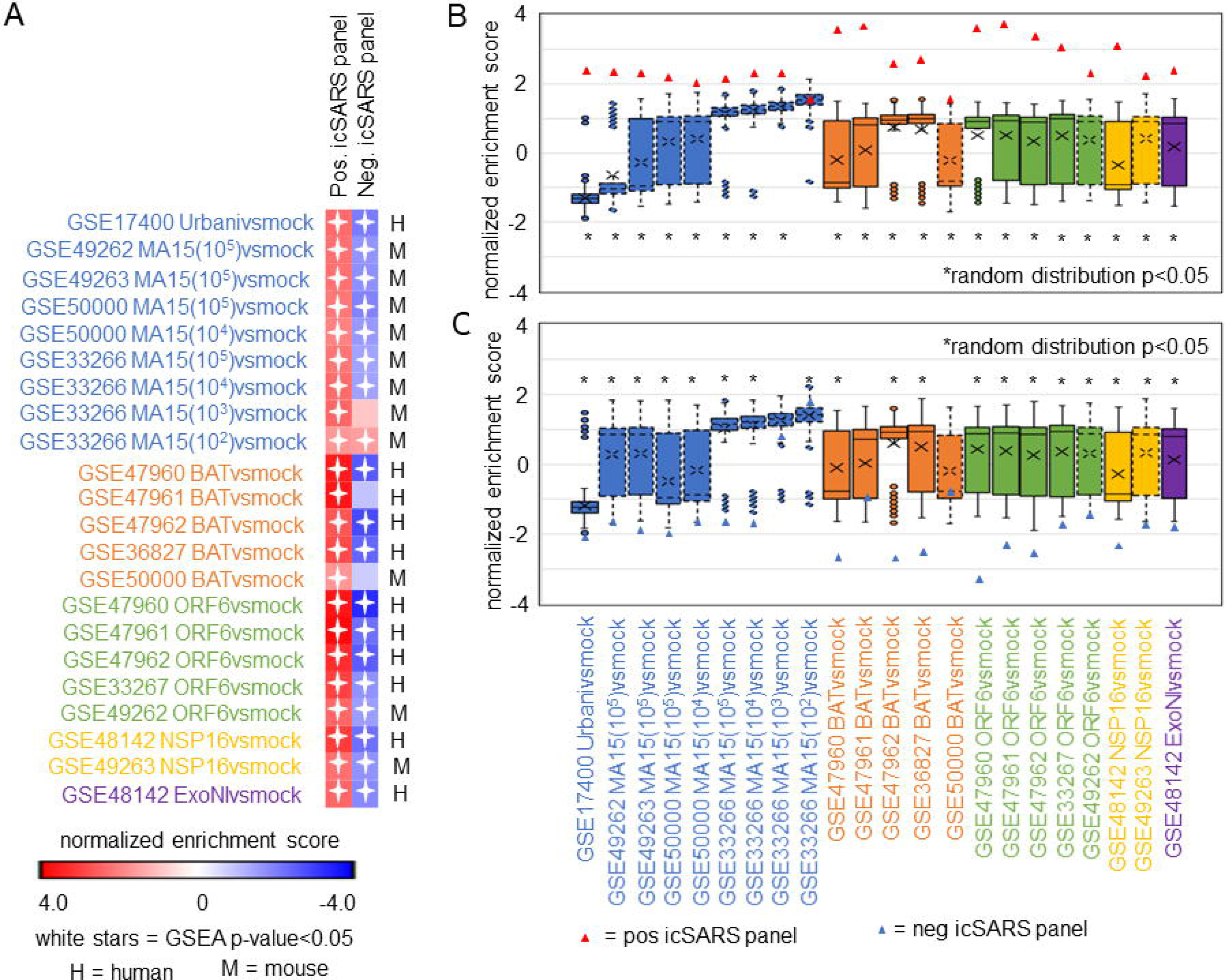
icSARS Panel Enrichment Detected Differential Gene Expression Similarities Across SARS Strains with Varying Virulen A) Heat map of Gene Set Enrichment Analysis (GSEA) calculated normalized enrichment scores (NES) of the positive and negative icSARS panels across SARS-CoV strains with varying levels of virulence in both human lung cultures and mouse lung samples. B) Box and whisker plots of NES from 1000 randomly generated gene panels containing 233 genes (individual queries) compared to gene signatures (individual references) used in (A). C) Box and whisker plots of NES from 1000 randomly generated gene panels containing 114 genes (individual queries) compared to gene signatures (individual references) used in (A).

### 3.5 Meta-analysis of Leading-edge Genes from Positive icSARS Panel GSEA Revealed Five Top Gene Candidates

To determine which positive icSARS panel genes were most associated with SARS-CoV infections, we compared inclusion of leading-edge genes identified by GSEA across 22 gene signatures derived from five SARS-CoV strains. We examined leading-edge gene inclusion in each SARS-CoV strain specifically (Figure 5A). Genes identified through leading-edge intersections represent genes associated with infection of that specific SARS-CoV strain. We noted 13 of the 61 leading-edge genes identified from the Urbanivsmock signature in human lung cultures were shared in leading-edges across the seven MA15vsmock signatures from mouse lung samples (Supplemental Material STables 7-8). There were 18 leading-edge genes from the Urbanivsmock signature that were not included across all platforms used to profile MA15 gene expressions and 10 positive icSARS panel genes not included in all platforms used to profile Urbani and MA15 gene expressions. We found 12 of the 49 leading-edge genes identified across the four BATvsmock signatures from human lung cultures were shared in BATvsmock signature leading-edge from mouse lung samples (Supplemental Material STables 9-10). There were 20 positive icSARS panel genes not included in all platforms used to profile BAT gene expressions that we included in further analysis with 10 of these genes shared across leading-edges from BATvsmock human culture signatures. Out of the 49 leading-edge genes identified across the four ORF6vsmock signatures from human lung cultures, 12 were shared with the ORF6vsmock signature leading-edge from mouse lung samples (Supplemental Material STables 11-12). There were 14 positive icSARS panel genes not included in all platforms used to profile ORF6 gene expressions. Finally, we found 43 of the 99 leading-edge genes identified in the NSP16vsmock signature from human lung cultures were shared in the NSP16vsmock signature leading-edge from mouse lung samples (Supplemental Material STables 13-14). There were 21 positive icSARS panel genes not included in all platforms used to profile NSP16 gene expressions. There were 73 leading-edge genes identified from the one ExoNIvsmock gene signature used in this study (Supplemental Material STable 15). Overall, we identified several genes with known associations to infection with specific SARS-CoV strains, such as *fos* and *jun* that were found in leading-edges from signatures derived from dORF6 [2] and Urbani [32] infections in human lung cell cultures. We also found several genes with no previously reported associations to that specific SARS infection. These findings were akin to findings from our meta-analysis of icSARS infection earlier that demonstrate the ability of our meta-analysis approach to identify genes associated with specific SARS infections and predict new genes associated with infections of specific SARS strains.

**Figure 5.**
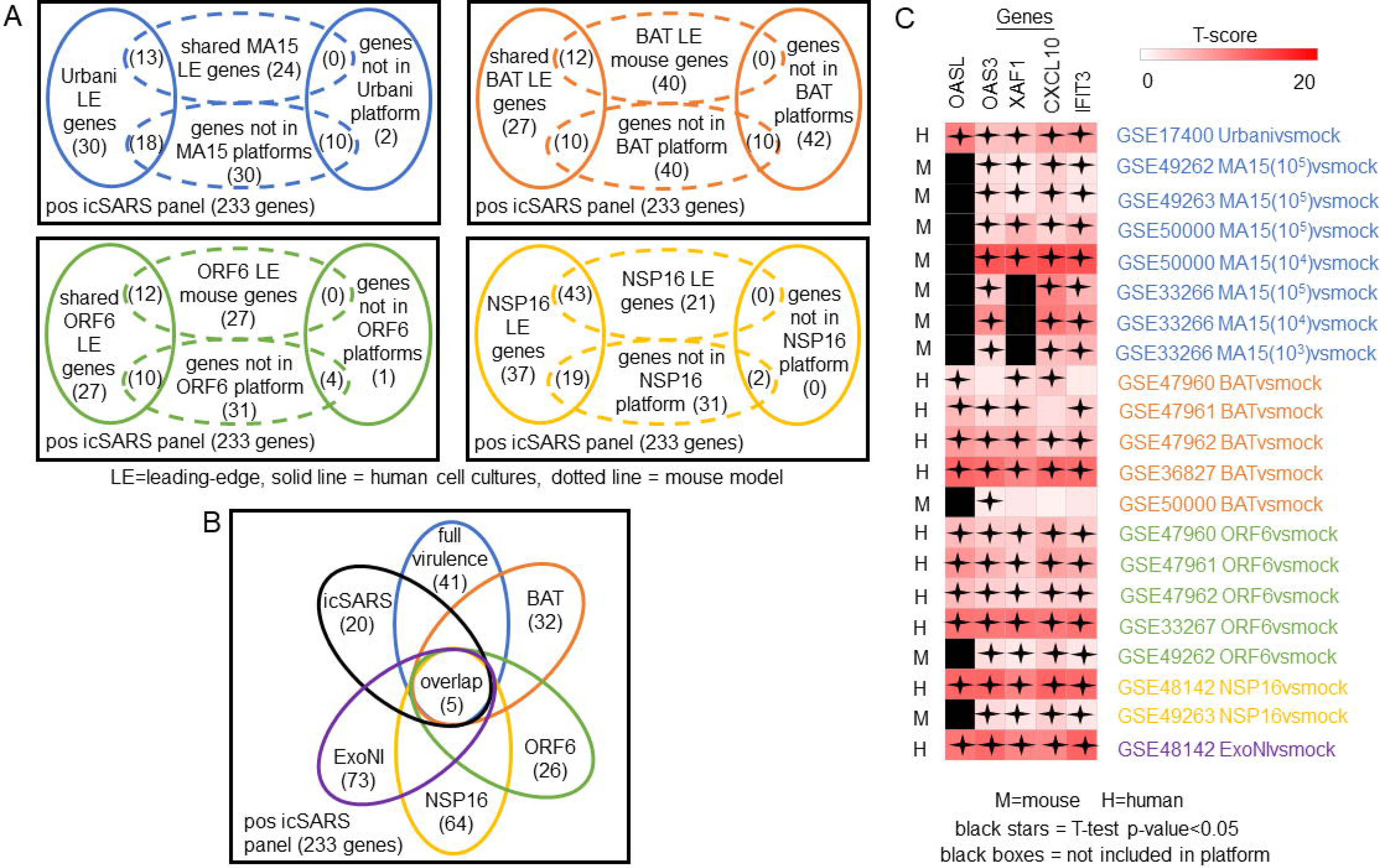
Meta-analysis Across 28 Gene Signatures Representing Seven SARS-CoV Strains Varying in Virulence Identified Five Over-Expressed Genes Associated with SARS-CoV Infection A) Venn diagrams of the inclusion and overlap of positive icSARS panel genes in identified leading-edges and dataset platforms from human and mouse gene signatures shared in individual strains of SARS-CoV. B) Venn diagram of the inclusion and overlap of shared positive icSARS panel genes identified in SARS-CoV leading-edges across individual strains of SARS-CoV. C) Heat map of T-scores for the five positive icSARS panel leading-edge genes identified in (B).

Next, we analyzed the intersection of common leading-edge genes across all six SARS strains examined in this study. We found five positive icSARS panel genes, IFN-induced protein with C-X-C motif chemokine ligand 10 (CXCL10), 2′-5′-oligoadenylate synthetase 3 (OAS3), 2′-5′-oligoadenylate synthetase-like (OASL), tetratricopeptide repeats 3 (IFIT3), and XIAP associated factor 1 (XAF1), in all 22 positive icSARS panel leading-edges (Figure 5B). As additional verification of our findings, we re-ran GSEA using the panels generated from GSE47961-derived icSARSvsmock (500-gene queries) and noticed this change did not affect the inclusion of our five gene candidates across leading-edges. We further noted all five gene candidates identified here were included in leading-edges identified by the 500 gene query set size at the 72hr and 96hr time points first examined in Supplemental Material SFig 1, suggesting that these gene candidates likely would have been identified if later time points were selected for use, thus not altering our overall result. Differential gene expression (*i.e.*, T-scores) heat maps illustrated the strong consistency and extent of expression changes observed across gene signatures (Figures 3C and 5C), further supporting the conclusion that these five genes were associated with SARS-CoV infection regardless of strain.

### 3.6 Top Five Gene Candidates Also Associated with MERS-CoV and SARS-CoV2

We expanded our analysis to examine icSARS panel enrichment and inclusion of leading-edge genes identified by GSEA across five MERS-CoV gene signatures and three SARS-CoV2 signatures (Figure 6). Two MERS-CoV gene signatures and all SARS-CoV2 signatures were in human lung cultures while the other three MERS-CoV gene signatures were derived from mouse lung cultures with various inoculation doses [55]. Using GSEA to calculate enrichment between icSARS panels and MERS-CoV, we found both icSARS panels significantly enriched (NES>1.97, GSEA p-value<0.001 for positive panel, NES<-1.68, p-value<0.003 for negative panel) across all five gene signatures (Figure 6A). For comparison to SARS-CoV2 gene signatures, we found the positive icSARS panel significantly enriched (NES>1.85, p-value<0.001) while the negative icSARS panel was not consistently enriched (NES<−1.13, p-value<0.210), which was not surprising since enrichment of the negative icSARS panel was inconsistent in icSARS signatures. We confirmed that achieved significant NES for the positive icSARS panel were non-random (null distribution p-value<0.02, Figure 6B) via random modelling for all signatures, which did not hold true for the negative icSARS panel (Figure 6C, p-value<0.573). When examining leading-edge gene inclusion in each SARS strain specifically (Figure 6D), we noted 32 of the 60 leading-edge genes identified from two MERSvsmock signatures in human lung cultures were shared in leading-edges across the three MERSvsmock signatures from mouse lung cultures (Supplemental Material STables 16-17). There were six leading-edge genes from MERSvsmock signatures in human cultures that were not included across all platforms used to profile MERS gene expressions. For SARS-CoV2 signatures, there were 29 leading-edge genes shared across the three signatures (Supplemental Material STable 18). When looking at leading-edges across all SARS strains examined in this study, we again found the same five gene candidates with strong consistency and extent of expression changes observed across gene signatures as determined by T-score (Figure 6E), supporting the conclusion that these five genes were associated with SARS infection overall.

**Figure 6.**
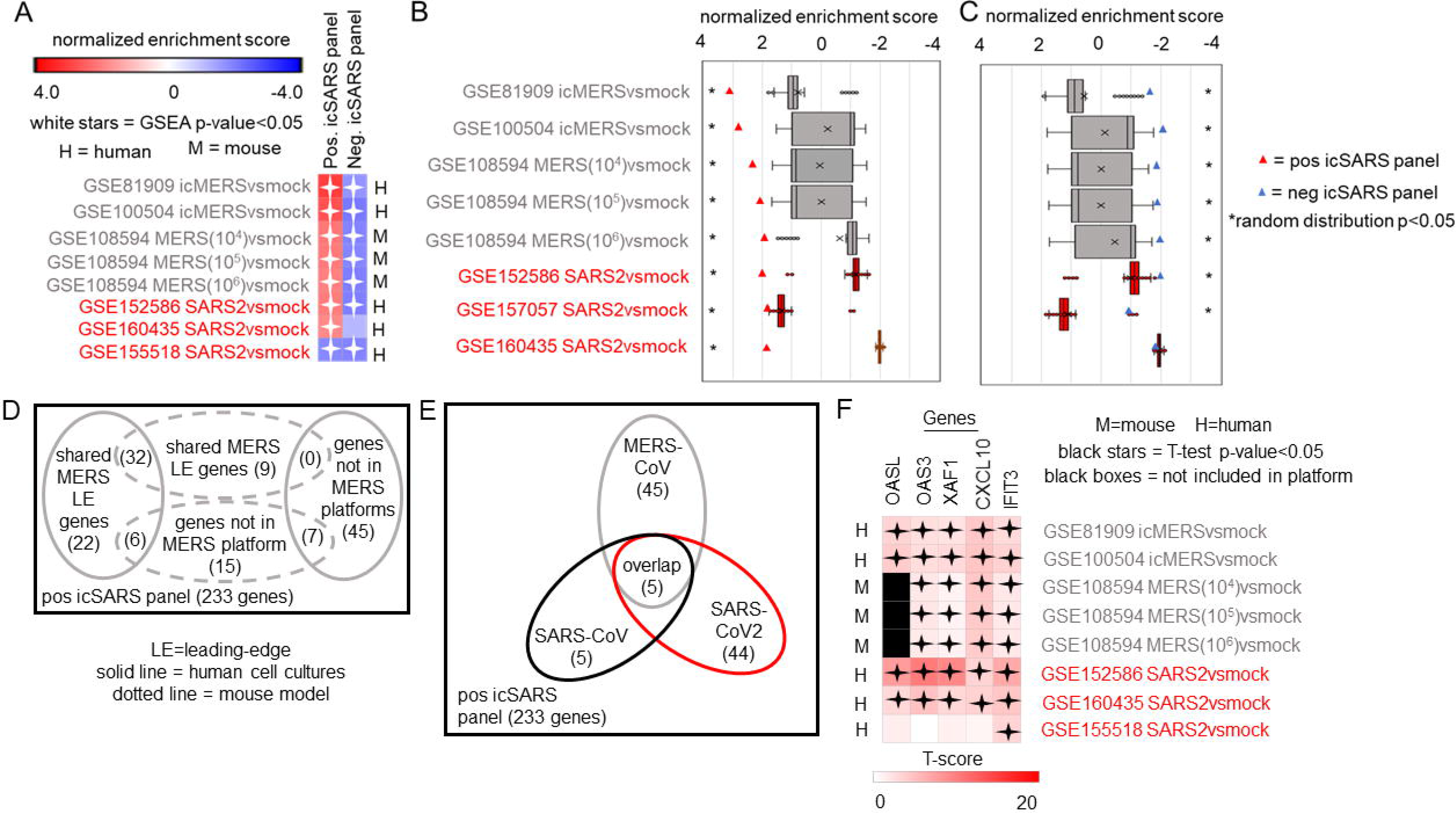
Five Over-Expressed Genes Identified in SARS-CoV Meta-analysis Found in Meta-analysis of MERS-CoV and SARS-CoV2 Signatures A) Heat map of Gene Set Enrichment Analysis calculated normalized enrichment scores for positive and negative icSARS panels across gene signatures derived from MERS-CoV and SARS-CoV2 infections in human or mouse lung cultures. B) Box and whisker plots of normalized enrichment scores from 1000 randomly generated gene panels containing 233 genes (individual queries) compared to MERS-CoV and SARS-CoV2 gene signatures (individual references). C) Box and whisker plots of normalized enrichment scores from 1000 randomly generated gene panels containing 114 genes (individual queries) compared to MERS-CoV and SARS-CoV2 gene signatures (individual references). D) Venn diagram of the inclusion and overlap of positive icSARS panel genes in identified leading-edges and dataset platforms across MERS-CoV human and mouse gene signatures. E) Venn diagram of the inclusion and overlap of shared positive icSARS panel genes in identified in SARS-CoV (Figure 6), MERS-CoV (from D), and SARS-CoV2 gene signatures. F) Heat map of T-scores for the five positive icSARS panel leading-edge genes identified in (E).

## 4 Discussion

SARS infections remain a serious public health threat due to their strong pandemic causing potential. While efforts have gone into developing effective therapeutics for SARS-infected patients, treatment options were still limited, due in part to an incomplete understanding of the molecular changes driving SARS infections. Identification of differentially expressed genes associated with SARS infections can improve our understanding of SARS-induced molecular changes, potentially contributing to the development of new therapeutic options to use in the fight against SARS and future SARS outbreaks. This work performed a meta-analysis of gene signatures generated from mRNA expression data across SARS-CoV, MERS-CoV, and SARS-CoV2 infections to reveal differentially expressed genes associated with SARS infections.

Among genes identified by our GSEA-based meta-analysis approach, we found five IFN-inducible gene candidates, CXCL10, OAS3, OASL, IFIT3, and XAF1, stood out consistently across SARS strains. CXCL10 was an IFN γ-induced protein with a strong connection to inflammatory and infectious diseases including viral infections [63]. CXCL10 was associated with infections of several SARS strains, specifically icSARS [30] and Urbani [32] in human lung cell cultures, MA15 in mouse lung samples [35], and SARS-CoV2 in clinical bronchoalveolar lavage fluid and plasma samples [64]. Also, increased levels of CXCL10 were associated with acute respiratory distress syndrome (ARDS), a clinical result of the cytokine storm frequently described across SARS infections, especially SARS-CoV2 where high CXCL10 levels have been implicated in increased disease severity and poorer patient outcomes [24, 64–67]. The two OAS genes (OAS3 and OASL) were anti-viral restriction factors [68]. Both OAS genes had reported associations with Urbani infections in human lung cell cultures [32]. Further, associations between the OAS pathway and MERS-CoV and SARS-CoV2 infections have been reported [69] and single nucleotide polymorphisms in OAS genes were found to be involved in the protective effects of Neandertal haplotypes against SARS-CoV2 [70, 71]. IFIT3 was one of four IFN-induced proteins with tetratricopeptide repeats whose expression was greatly enhanced by viral infection, IFN treatment, and pathogen-associated molecular patterns [72]. While IFIT3 was not specifically named in published SARS-CoV reports examining these datasets used in this study, IFIT1 had reported associations with MA15 infections in mouse lung samples [34, 35]. All IFIT proteins including IFIT3 have been connected to SARS-CoV2 infection in human lung cell cultures and samples [73]. XAF1 was an apoptotic gene whose reduced or absent expression in tumor samples and cell lines leads to poorer survival in gastric adenocarcinomas [74, 75]. This paper was the first study to report an association between XAF1 and SARS-CoV infections for any strain, to the best of our knowledge, though a recent report examining differential gene expression associated with SARS-CoV-2 infection identified increased expression of CXCL10, OAS3, IFIT3, and XAF1 in human epithelial lung cells and lung samples from *Cynomolgus maca* (cynomolgus monkey) and mice [36]. Taken together, these results supported the conclusion that targeting IFN response therapeutically, particularly one or more of these five identified genes, might improve outcomes for patients with SARS infections. Our results supported recent reports detailing the connections between SARS-CoV2 and type I IFN response, including one report demonstrating that pretreatment with IFNβ protected both Calu3 and Caco2 (human intestinal epithelium) cells against SARS-CoV2 infection [26, 73]. Several reports also examine the success of IFN therapy for SARS-CoV2 patients with promising results though questions around timing, type of IFN, and administration route remain debatable [21, 25–27, 76]. While these five genes were noted among those identified by pathway enrichment analysis like the one performed here using GO (Supplemental Material STable 3), our meta-analysis approach improves upon the existing method by refining the gene candidate list and examining a larger number of datasets.

In our study, we observed our meta-analysis approach had threshold of gene detection limits that may have biological implications. For example, we failed to observe non-random enrichment in the GSE33266 MA15(10^2^)vsmock signature, which was the lowest inoculation dose used in this meta-analysis (Figure 4B). While we removed the GSE33266 MA15(10^2^)vsmock signature from inclusion in our meta-analysis because its observed enrichment was not statistically different from NES achieved randomly, we noted that IFIT3 was the only top candidate in the leading-edge of the MA15(10^2^)vsmock signature and IFIT3 was not statistically significant individually (Welch’s T-test p-value=0.160). These findings suggest there was a lower inoculation dose limit to our approach’s ability to detect relevant genes, which should be considered when applying our meta-analysis to future research. Further, since gene expression has been known to change over a time course [2], we repeated the process of defining icSARS gene panels over a range of time points (24, 48, 72, and 96hrs) and found consistent enrichment and identification of gene candidates for all timepoints but the 24hr timepoint (Supplemental Material SFig 1). This finding suggested that gene expression changes generated by an icSARS infection become more predictable at later time points. These gene detection threshold limits were interesting considering recent reports of that SARS-CoV2 suppresses IFN response in early infection stages to establish infection and early IFN treatments of SARS-CoV2 patients improved mortality [21, 27, 73]. We cannot rule out the idea that the threshold of gene detection limits observed in our study were not the result of IFN therapeutic limitations previously reported.

While this meta-analysis revealed genes with already well-established associations to SARS infections, this purely bioinformatic work was limited by a lack of direct experimental evidence. We were unable to conduct follow-up experiments using other techniques, such as Western blotting or qRT-PCR, to confirm our top gene candidate predictions that were generated only from mRNA expression data. Further, datasets selected for this study contained only human lung cultures or mouse lung samples due to their public availability at the time this study was conducted and inclusion of 48hr time point data. Based on results from this study, gene expression data from IFN-treated and untreated SARS-CoV2 patients would be of particular interest for future studies. We predict IFNtreatedSARS2vsuntreatedSARS2 gene signatures would be reversed (*i.e.*, positive icSARS panel achieves significant enrichment with a negative NES), indicating IFN-treated samples looked more like untreated cultures and samples from this study, further supporting the conclusion of targeting IFN as viable therapeutic option for SARS infections.

## 5 Conclusion

This work used mRNA expression data to predict genes associated with SARS infections through a meta-analysis examining gene signatures. Through our GSEA-based meta-analysis approach, we identified five over-expressed IFN-inducible genes as being most associated with SARS infections. We concluded from this finding that targeting type I IFN response either in a stand-alone or combination therapy, particularly the five genes identified here, might improve treatment options for SARS and other β-coronavirus infections. Our conclusion supported prior reports of successful outcomes from IFN therapy in patients with less severe SARS infections and at earlier time points. Overall, this work demonstrated the gene detection ability and reproducibility of our meta-analysis approach and presents it as a useful computational approach through application on mRNA expression data from SARS and mock infected cultures and samples.

## 6 Declarations

### 6.1 Ethics approval and consent to participate

Manuscript does not generate new data from human participants or animal models. Not applicable.

### 6.2 Consent for publication

Manuscript does not contain data from any individual person. Not applicable.

### 6.3 Availability of data and materials

The datasets analyzed during the current study are available in the Gene Expression Omnibus repository https://www.ncbi.nlm.nih.gov/geo/.

### 6.4 Competing interests

The authors declare that the research was conducted in the absence of any commercial or financial relationships that could be construed as a potential conflict of interest.

### 6.5 Funding

No funding support was provided for this work.

### 6.6 Authors’ Contributions

L.H. lead manuscript preparation. A.P. generated the supplemental materials and time course data. All authors contributed to the interpretation of results and manuscript review.

## 6.7 Acknowledgments

The authors would like to thank Jared Geller for the pathway analysis and Terri Pulice for the graphic design assistance.

## Notes

### Competing Interest Statement

The authors have declared no competing interest.

### Summary of Updates

Additional analysis with MERS and SARS-CoV2, expanded introduction, revised conclusions.

