## Supplemental Materials Figs for "Gene Expression Meta-Analysis Reveals Interferon-induced Genes"

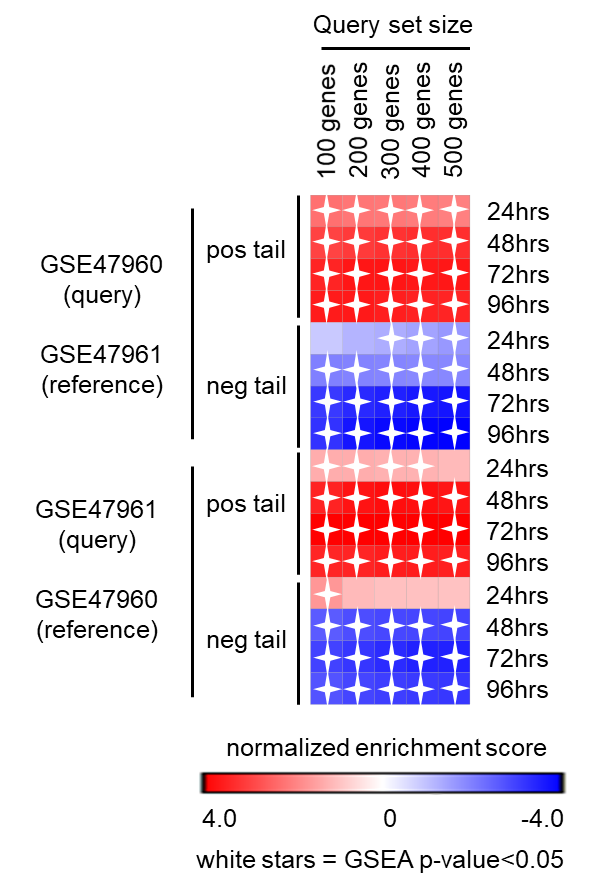

**Supplementary Figure 1.** Heat map of Gene Set Enrichment Analysis (GSEA) calculated normalized enrichment scores showed consistent significant enrichment for various sized query gene sets generated from either positive and negative tails of GSE47960- or GSE47961-derived icSARSvsmock signatures at 48, 72, and 96hrs, but not at the 24hr timepoint.

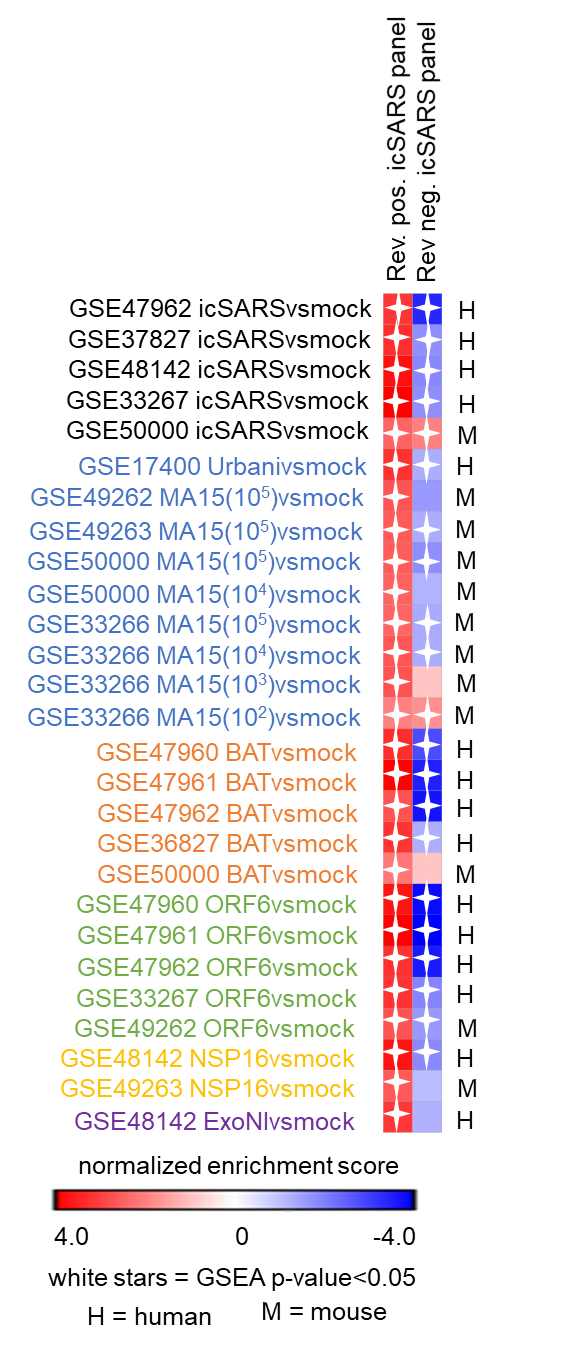

**Supplementary Figure 2.** Heat map of Gene Set Enrichment Analysis (GSEA) calculated normalized enrichment scores show consistent significant enrichment of the reverse (*i.e.*, derived from GSE47960 as reference and GSE47961 to define query gene sets) positive icSARS panel across SARS-CoV strains with varying levels of virulence in both human lung cultures and mouse lung samples. Enrichment for the reverse negative icSARS panel was inconsistent across SARS strains. These results were like those achieved from positive and negative icSARS panels (*i.e.*, derived from GSE47961 as reference and GSE47960 to define query gene sets, Figure 4).
